# Induction of specific antibodies, IgG-secreting plasmablasts and memory B cells following BCG vaccination

**DOI:** 10.1101/2021.02.18.431837

**Authors:** Julia Bitencourt, Morven Wilkie, Marco Polo Peralta Alvarez, Ashley Jacobs, Daniel Wright, Stephanie A. Harris, Steven G. Smith, Sean Elias, Andrew White, Sally Sharpe, Matthew K. O’Shea, Helen McShane, Rachel Tanner

**Affiliations:** The Jenner Institute, University of Oxford, UK; Gonçalo Moniz Institute, Oswaldo Cruz Foundation (FIOCRUZ), Salvador, Brazil; University of Cape Town, Cape Town, South Africa; London School of Hygiene and Tropical Medicine, UK; Division of Biosciences, Brunel University, UK; Public Health England, Porton Down, Salisbury, UK; Institute of Immunology and Immunotherapy, College of Medical and Dental Sciences, University of Birmingham, Birmingham, UK

**Keywords:** humoral, antibody, B cells, tuberculosis, BCG, vaccine

## Abstract

Many tuberculosis (TB) vaccine candidates are designed as a boost to BCG; an understanding of the BCG-induced immune response is therefore critical, and the opportunity to relate this to circumstances where BCG does protect may direct the design of more efficacious vaccines. While the T cell response to BCG vaccination has been well-characterised, little is known about the B cell and antibody response. We demonstrate BCG vaccine-mediated induction of specific antibodies in different human populations and macaque species which represent important preclinical models for TB vaccine development. We observe a strong correlation between antibody titres in serum versus plasma with modestly higher titres in serum. We also report for the first time the rapid and transient induction of antibody-secreting plasmablasts following BCG vaccination, together with a robust and durable memory B cell response in humans. Finally, we demonstrate a potential contribution of the antibody response to BCG vaccine-mediated control of mycobacterial growth *in vitro*. Taken together, our findings indicate that the humoral immune response in the context of BCG vaccination merits further attention to determine whether TB vaccine candidates could benefit from the induction of humoral as well as cellular immunity.

## 1.0 Introduction

Tuberculosis (TB), caused by *Mycobacterium tuberculosis* (*M.tb*), remains a major global health threat with 10 million new cases and 1.4 million deaths in 2019 [1]. There are currently no validated correlates of protection from TB. However, *M.tb* is an intracellular pathogen and the necessity for a T cell response in conferring acquired immunity to TB has been demonstrated in numerous studies [2–6]. This has led to a limited focus on the humoral response to TB, although emerging evidence suggests that antibodies may play a more significant role in protection than previously appreciated [7–10]. Antibodies could contribute to protection directly through increasing phagocytosis and phagolysosome formation or bacterial neutralisation, and/or indirectly through enhancing T cell-mediated immunity [7]. It has recently been shown that compared to antibodies from patients with active TB disease (ATB), antibodies from individuals with latent TB infection (LTBI) have unique Fc functional profiles, selective binding to FcγRIII and distinct antibody glycosylation patterns, and also that they drive enhanced phagolysosomal maturation, inflammasome activation and macrophage killing of intracellular *M.tb* [9].

Bacillus Calmette Guérin (BCG) is the only currently available vaccine against TB. BCG confers incomplete and variable protection against pulmonary TB in adolescents and adults, and a new more efficacious TB vaccine is needed [11, 12]. However, it is uncertain which aspects of the immune response a candidate vaccine should aim to induce in order to confer protection that is superior to BCG. Due to the important role of BCG vaccination in protecting infants from severe forms of TB disease, and its potential non-specific effects protecting from all-cause mortality, most TB vaccine candidates are designed as a heterologous boost to a BCG prime [13, 14]. It is thus critical to understand the immune response to BCG vaccination and which aspects will be, or would ideally be, induced or boosted. Furthermore, because BCG vaccination is partially protective, and can confer superior protection when administered intravenously in macaques [15, 16] or as a revaccination in South African adolescents [17], studying the immune response to BCG offers a valuable opportunity to explore immune mechanisms of protection and apply these to inform the design of more efficacious TB vaccine candidates.

Robust Th1 responses to BCG vaccination have been described and are generally considered to be essential, but not sufficient, for protection [18–21]. A role for trained innate immunity, unconventional T cells and humoral immunity in BCG-mediated protection has been proposed [22–24]. Indeed, in a *post-hoc* correlates of risk analysis, levels of Ag85A-specific IgG were associated with reduced risk of TB disease in BCG-vaccinated South African infants [25]. We have recently comprehensively reviewed what is known about the humoral immune response to BCG vaccination and revaccination across species [26]. In brief, the literature presents inconsistent evidence for the induction of specific antibody responses following BCG vaccination, and the relevance of these responses is unclear, with support both for [25, 27–30] and against [28, 31–33] a protective function.

Antibodies are produced by antibody-secreting B cells (ASCs). Upon activation by recognition of their cognate antigen during infection or vaccination, B cells undergo clonal expansion and differentiate into plasmablasts and memory B cells (mBCs) [34]. While plasmablasts and plasma cells secrete antibody, mBCs can survive quiescently for decades, poised to rapidly respond to antigen restimulation by differentiating into short- and long-lived ASCs that can prolong the duration of high serum antibody levels [35, 36]. It is generally accepted that mBC-derived ASCs are central to the long-term protection mediated by most licenced vaccines [34, 37], and the dynamics, magnitude and specificity of the ASC response to vaccination against other infections have been described [38–40]. While one study has reported a significantly higher frequency of purified protein derivative (PPD)-specific mBCs in peripheral blood mononuclear cells (PBMC) from historically BCG-vaccinated compared with unvaccinated volunteers [41], the longitudinal ASC response following BCG vaccination has not been described. Evaluating these cells may provide a more dynamic, discriminative measure of humoral immunity than detecting stable serum antibodies by conventional serology.

The hitherto poorly-defined humoral response to BCG vaccination merits further investigation. We set out to explore the BCG vaccine-induced specific antibody response in serum and plasma collected from different cohorts of human volunteers representing TB-endemic and non-endemic populations. As the macaque model is widely used in TB studies [42], we included macaque samples to better understand the translatability of antibody responses to humans. We also sought to determine the frequency and kinetic of BCG-induced antigen-specific ASCs (both plasmablasts and mBCs), and their association with serum antibody levels. Finally, we employed the direct PBMC mycobacterial growth inhibition assay (MGIA) to explore a potential functional contribution of BCG-induced antibodies to the control of mycobacterial growth. Our findings offer proof-of-concept and provide some logistical foundations for a more comprehensive characterisation of the BCG vaccine-induced antibody response across different populations and vaccine regimens for which different levels of protection are observed.

## 2.0 Methods

### 2.1 Human samples

Human PBMC, serum and plasma samples were taken from a randomised controlled clinical study of healthy BCG-naïve UK adults, henceforth referred to as Study 1. Volunteers were randomised to receive BCG SSI at a standard dose (2-8 x 10^5^ CFU) (n=27) or to be unvaccinated controls (n=8). There was a similar age range between groups (mean 27 years, range 18-42 for BCG vaccinees; mean 30 years, range 20-45 for controls). 74% of vaccinees and 50% of controls were female. Samples were used from baseline and follow-up visits at 7, 14, 21, 28 and 84 days. ELISA operators were blinded to treatment group. The study was approved by the NHS Research Ethics Service (NRES) Committee South Central - Oxford B (REC reference 15/SC/0022), and registered with ClinicalTrials.gov (NCT02380508). All aspects of the study were conducted according to the principles of the Declaration of Helsinki and Good Clinical Practice. Volunteers provided written informed consent prior to screening. Baseline biochemical and haematological analysis and serological testing for HIV, HBV and HCV were performed to ensure no abnormalities warranting exclusion. LTBI was excluded at screening by T Spot.TB (Oxford Immunotec, UK) or QuantiFERON^®^-TB Gold In-Tube test.

PBMC and serum samples from Study 2 were collected from a previously-described cohort of adult male military recruits who had recently arrived in the UK from Nepal [43]. Volunteers shown to be LTBI-negative by T-SPOT.TB ELISpot assay (Oxford Immunotec, UK) and without known history or physical evidence of prior BCG vaccination were enrolled into the current study (group A, n=11), together with a small group of historically BCG-vaccinated individuals who received no intervention (group B, n=4). All volunteers were male with a similar median age in groups A and B (18.9 vs. 20.5 years respectively). Ethical approval was granted by the Ministry of Defence Research Ethics Committee (MODREC 237/PPE/11), and all participants provided written informed consent. Group A received a single intradermal vaccination of BCG SSI at a standard dose (2-8 x 10^5^ CFU) in accordance with current British Army public health policy, and samples were collected at baseline and follow-up visits at 7 days and 70 days.

### 2.2 Macaque samples

Macaque serum and/or plasma samples used in the ELISA studies described here were collected as part of five independent historical studies of BCG vaccination at Public Health England (PHE), further details of which are provided in Table 1. Animals were randomised by socially-compatible group to receive an adult human dose of BCG Danish strain 1331 (SSI, Copenhagen) 2-8×10^5^ CFU intradermally (ID) into the upper arm under sedation, or to be unvaccinated controls. The BCG vaccine was prepared and administered according to the manufacturer’s instructions for preparation of vaccine for administration to human adults, by addition of 1ml Sauton’s diluent to a lyophilised vial. In all cases, animals were obtained from established breeding colonies at PHE in the UK, which were captive-bred for research purposes, and a single animal was considered an experimental unit. They were provided with enrichment in the form of food and non-food items on a daily basis; animal welfare was monitored daily. Study design and procedures were approved by the Public Health England Animal Welfare and Ethical Review Body and authorized under an appropriate UK Home Office project license. Animals were housed in compatible social groups and in accordance with the Home Office (UK) Code of Practice for the Housing and Care of Animals Used in Scientific Procedures (1989) and the National Centre for Refinement, Reduction and Replacement (NC3Rs) Guidelines on Primate Accommodation, Care and Use, August 2006 (NC3Rs, 2006).

**Table 1.** Summary of macaque samples used in antibody studies.

| Study | Species | Genotype | Age (yrs) | Sex | Groups (n) | Times | Ref. |
| --- | --- | --- | --- | --- | --- | --- | --- |
| 1 | Rhesus | Indian | 14-15 | 14 F<br>1 M | Naïve controls (5)<br>BCG vaccinated (10) | D0<br>D28<br>D56 | n/a |
| 2 | Rhesus | Indian | 2.9-3.2 | M | Naïve controls (4)<br>BCG vaccinated (5) | D0<br>D56 | [76] |
| 3 | Rhesus | Indian | 2.5 | M | BCG vaccinated (6) | D0<br>D56<br>D140 | [15] |
| 4 | Rhesus | Indian | 3.8-4.2 | M | BCG vaccinated (6) | D0<br>D56<br>D140 | [77] |
| 5 | Cynomolgus | Mauritian | 2.3-3.7 | M | BCG vaccinated (12) | D0<br>D56<br>D140 | n/a |

### 2.3 Enzyme-linked immunosorbant assay (ELISA)

ELISAs were performed as previously described [44]. The antigens used were purified protein derivative (PPD) from *M.tb* (Statens Serum Institut (SSI), Denmark) at a concentration of 5μg/ml; whole BCG SSI at a concentration of 5×10^5^ CFU/ml, *M.tb* H37Rv cell membrane fraction, culture filtrate or whole cell lysate all at a concentration of 2μg/ml; or lipoarabinomannan (LAM) at a concentration of 1μg/ml (BEI resources repository). Plates were coated with 50μl of antigen and incubated at 4°C overnight. Samples were prepared by diluting test serum and positive/negative control serum 1:10 (Figure 1), 1:50 (Figure 2) or 1:100 (Figure 4) and 50μl was added per well for 2 hours. For human samples, secondary antibody (goat anti-human γ-chain-specific whole IgG, α-chain-specific IgA, or μ-chain-specific IgM alkaline phosphatase conjugate, Sigma Aldrich, MO, US) was diluted 1:1000 and 50μl was added per well for 1 hour. For macaque samples, secondary antibody (goat anti-monkey γ-chain-specific whole IgG, α-chain-specific IgA, or μ-chain-specific IgM alkaline phosphatase conjugate, Rockland Immunochemicals, PA, US) was diluted 1:750 and 50μl added per well for 1 hour. 100μl of p-nitrophenyl phosphate (pNPP) development buffer was added to each well and the plates were read every 10 minutes using a Model 550 Microplate Reader (Bio-Rad, UK) until the positive control reached a predetermined OD_405_ that was consistent across plates. Reported values represent the mean OD_405_ values of triplicate negative controls subtracted from the mean OD_405_ of triplicate samples.

**Figure 1.**
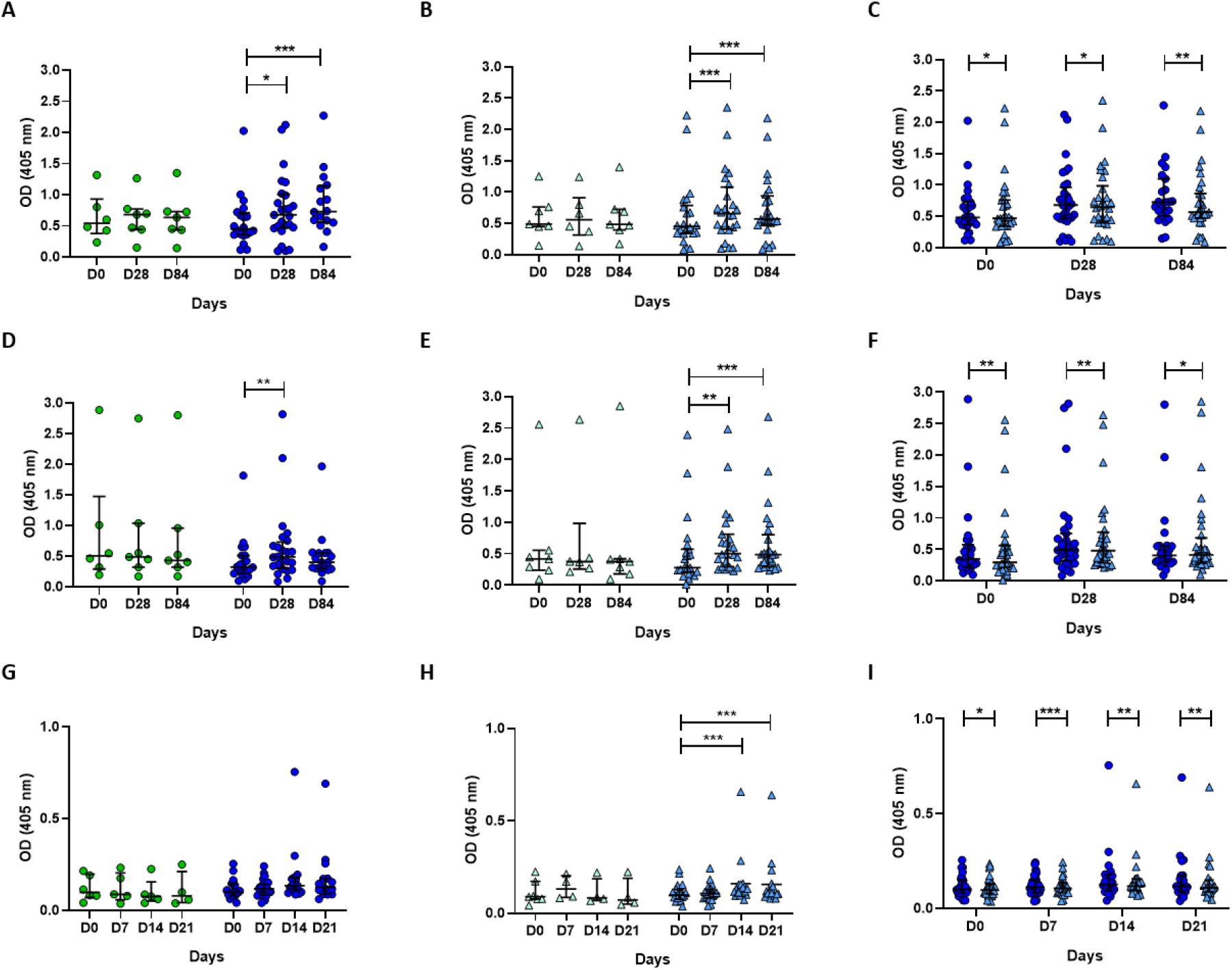
PPD-specific antibody responses to BCG vaccination in healthy UK adults. Serum (circles) and plasma (triangles) were collected from volunteers enrolled into human Study 1 who were either unvaccinated controls (green) or received BCG vaccination (blue). PPD-specific IgG (A-C), IgA (D-F) and IgM (G-I) responses were measured in serum (A, D, G) and plasma (B, E, H) over time, and directly compared between serum and plasma (C, F, I). Points represent the mean of triplicate values and bars show the median with the interquartile range (IQR). A mixed effects analysis with Dunnett’s correction for multiple comparisons was conducted on logged values to compare the BCG vaccine-induced response between post-vaccination and baseline time-points (A, B, D, E, G, H), and a Wilcoxon test was used to compare between serum and plasma responses at each time-point (C, F, I). * indicates a p-value of <0.05, ** indicates a p-value of <0.01, *** indicates a p-value of <0.001 and **** indicates a p-value of <0.0001.

**Figure 2.**
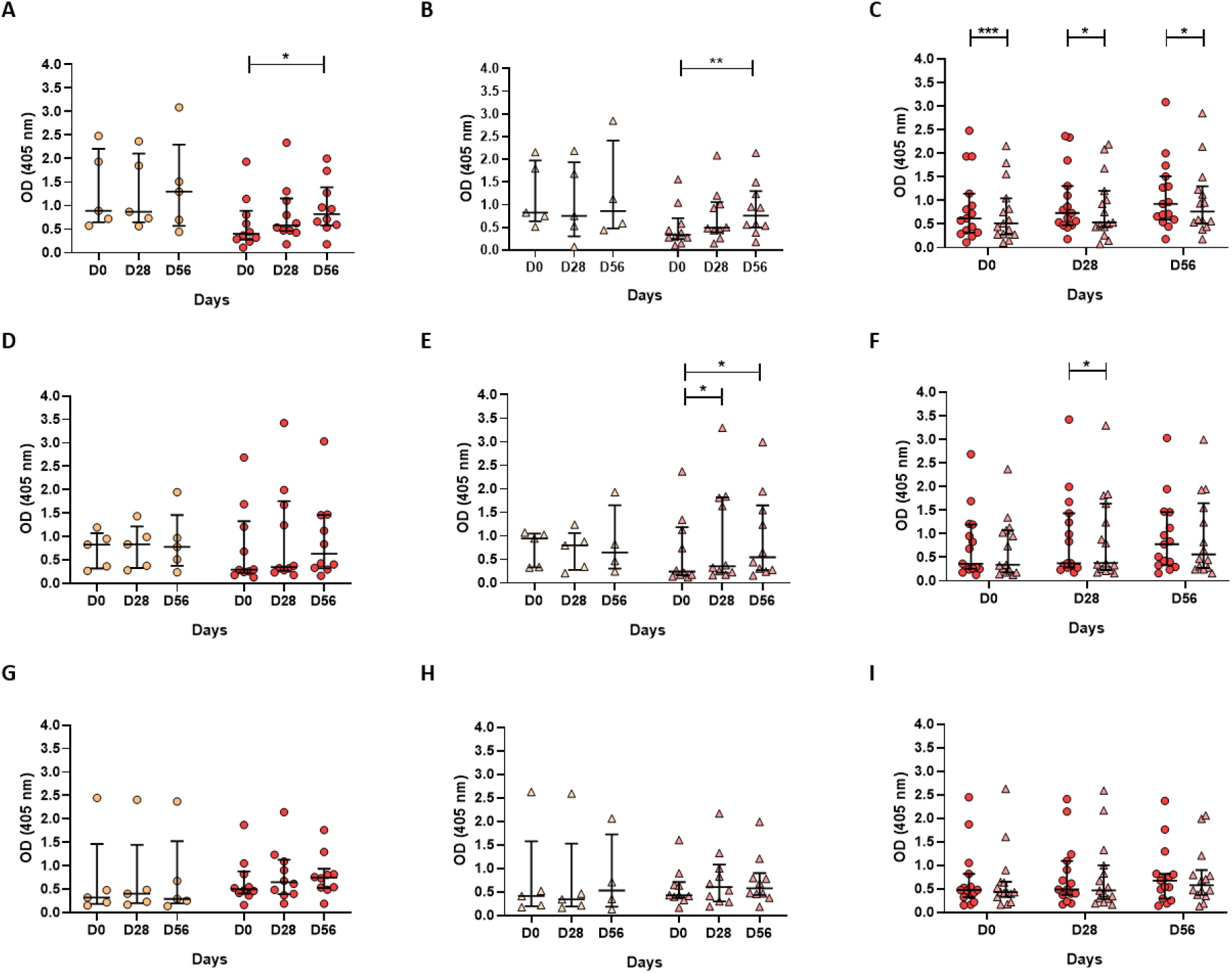
PPD-specific antibody responses to BCG vaccination in rhesus macaques. Serum (circles) and plasma (triangles) were collected from animals enrolled into macaque Study 1 which were either unvaccinated controls (orange) or received BCG vaccination (red). PPD-specific IgG (A-C), IgA (D-F) and IgM (G-I) responses were measured in serum (A, D, G) and plasma (B, E, H) over time, and directly compared between serum and plasma (C, F, I). Points represent the mean of triplicate values and bars show the median with the interquartile range (IQR). A Friedman test with Dunn’s correction for multiple comparisons was used to compare the BCG vaccine-induced response between post-vaccination and baseline time-points (A, B, D, E, G, H), and a Wilcoxon test was used to compare between serum and plasma responses at each time-point (C, F, I). * indicates a p-value of <0.05, ** indicates a p-value of <0.01, and *** indicates a p-value of <0.001.

**Figure 3.**
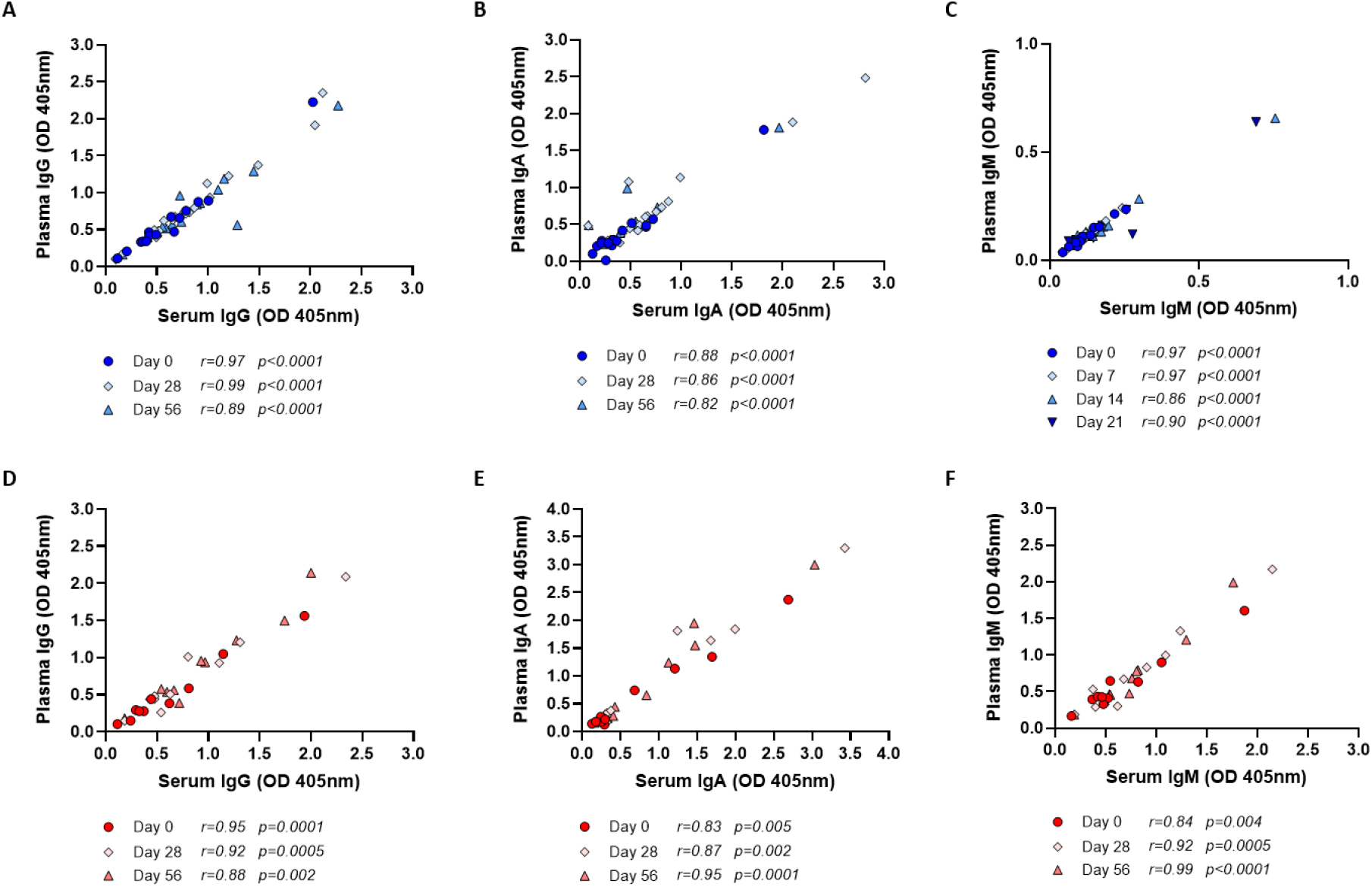
PPD-specific antibody responses to BCG vaccination compared between plasma and serum. Antibody responses in serum and plasma collected from healthy UK adults enrolled into human Study 1 (blue, A-C) and rhesus macaques enrolled into macaque Study 1 (red, D-F). IgG (A), IgA (B) and IgM (C) were compared at baseline and at 28 and 84 days post-BCG vaccination in humans. IgG (D), IgA (E) and IgM (F) responses were compared at baseline and at 28 and 56 days post-BCG vaccination in macaques. Spearman’s rank correlation was used in all cases.

**Figure 4.**
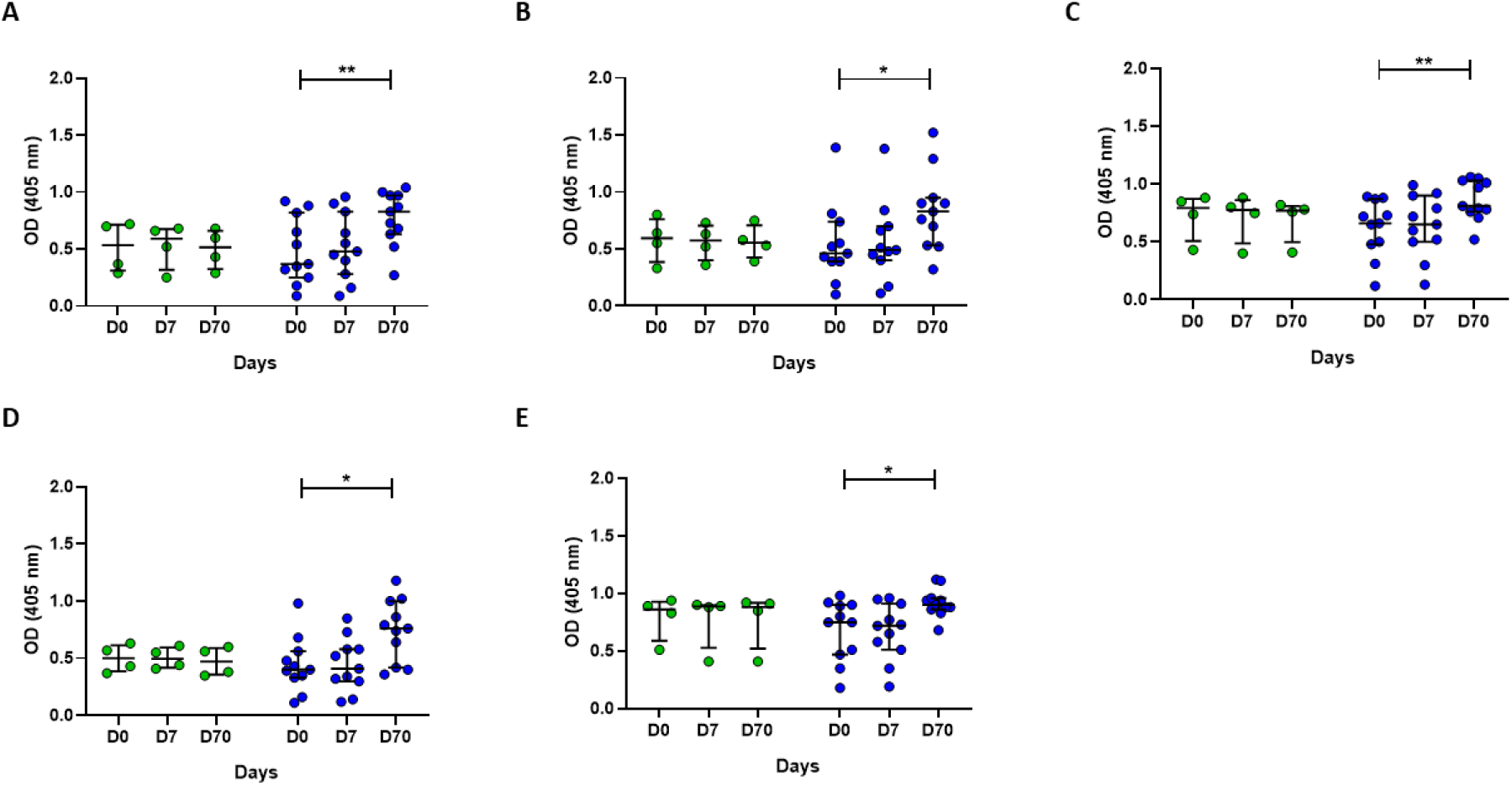
IgG responses to different mycobacterial fractions in serum collected from Nepalese military recruits. Serum samples were taken from Nepalese adults enrolled into human Study 2 (green = historically BCG-vaccinated individuals who received no intervention, blue = BCG vaccinees). IgG responses to whole BCG (A), M.tb whole cell lysate (B), M.tb cell membrane fraction (C), M.tb culture filtrate (D) or M.tb lipoarabaninomannan (E) were measured. Points represent the mean of triplicate values and bars show the median with the interquartile range (IQR). A repeated measures one-way ANOVA with Dunnett’s multiple comparisons test was used to compare between time-points in the BCG vaccinated group where * indicates a p-value of <0.05 and ** indicates a p-value of <0.01.

### 2.4 *Ex vivo* B cell enzyme-linked immunospot (ELISpot) assays

#### 2.4.1 Antibody-secreting cell (ASC) ELISpot

ELISpot plates were coated in triplicate with 50μl/well of BCG SSI at 4×10^6^ CFU/ml. Sterile PBS was used for negative controls. For the detection of total IgG-secreting cells, six wells were coated with 50μl/well of polyvalent goat anti-human IgG at 50μg/ml in PBS. Plates were incubated overnight at 4°C and then washed three times with sterile PBS and blocked with R10 media (100μl/well) for 1 hour at 37°C. Cryopreserved PBMC was thawed and prepared as previously described and resuspended to a concentration of 5×10^6^ cells/ml in R10. 50μl of cell suspension was added to the negative control wells and to the first three antigen-coated wells per volunteer. Cells were diluted 1:2 in R10 for the second three antigen-coated wells per volunteer. PBMC dilutions for the total IgG controls were 1:1, 1:25 and 1:50 in R10; 50μl of the IgG control dilutions were added to duplicate wells. Plates were incubated at 37°C, 5% CO_2_ for 16-18 hours. Cells were discarded and plates washed six times with PBS-Tween. 50μl/well of anti-human IgG (gamma-chain specific) conjugated to ALP and diluted 1:5000 in PBS was added and plates incubated for 4 hours at room temperature. Plates were then washed 6 times with PBS-Tween. 50μl/well of developer was added for 3-5 minutes until spots developed. Plates were washed thoroughly and dried overnight before counting on an AID ELISpot reader. Frequencies of antigen-specific ASCs were calculated by subtracting the mean count of the negative control wells from the test antigen wells and correcting for the number of PBMC in the well. ASC frequency was reported as both the number of ASCs per 1×10^6^ PBMC and also as the percentage of total IgG-secreting cells. Responses were considered positive if the count was two or more spots in each replicate well, and the total number of spots in the antigen-coated wells twice that observed in the blank control wells.

#### 2.4.2 Memory B cell (mBC) ELISpot

For each volunteer, 500μl of thawed PBMC at a concentration of 2×10^6^ cells/ml in R10 media was added to six wells of a 24-well flat-bottom tissue culture plate. 500μl of Mitogen stimulation mix containing *Staphylococcus aureus* Cowan (SAC, 1:2400 dilution), CpG (5μg/ml) and pokeweed mitogen (PWM, 1:6000 dilution from a 1mg/ml stock), was added to five wells per volunteer. 500μl R10 was added to an unstimulated well as a control. Plates were incubated at 37°C, 5% CO_2_, for 6 days. On day 5, ELISpot plates were coated overnight at described above. On day 6, plates were washed and blocked as described. The cultured cells set up on day 0 for each volunteer were harvested by gentle resuspension and the stimulated cells and unstimulated cells were each pooled, washed twice by centrifugation at 700g for 5 minutes at room temperature, resuspended in R10 and counted. Cells were then resuspended at 2×10^6^ cells/ml in R10, and 100μl was added to the negative control wells and the first three antigen-coated wells. For the total IgG wells, dilutions of 1:1, 1:50 and 1:100 were added to each well in duplicate. 100μl of unstimulated cells were added to duplicate IgG wells. Plates were then incubated at 37°C, 5% CO_2_, for 16-18 hours and developed and counted as described above. Frequencies of mBC were shown as the number of BCG-specific mBC-derived ASC per million cultured PBMC and as the proportion of mBC-derived ASC of total IgG-secreting ASC.

### 2.4 Direct PBMC mycobacterial growth inhibition assay (MGIA)

MGIAs were performed as previously described [44], co-culturing 3×10^6^ PBMC and ~500 CFU BCG Pasteur in a volume of 480μl RPMI (containing 2mM l-glutamine and 25mM HEPES), plus 120μl autologous serum per well of a 48-well-plate for 96 hours at 37°C and 5% CO_2_. At the end of the culture period, co-cultures were added to 2ml screw-cap tubes and centrifuged at 12,000rpm for 10 minutes. During this time, 500μl sterile water was added to each well to lyse adherent monocytes. Supernatants were removed from the 2ml screw-cap tubes, and water from the corresponding well added to the pellet. Tubes were pulse vortexed and the full volume of lysate transferred to BACTEC MGIT tubes supplemented with PANTA antibiotics and OADC enrichment broth (Becton Dickinson, UK). Tubes were placed on the BACTEC 960 instrument (Becton Dickinson, UK) and incubated at 37°C until the detection of positivity by fluorescence. On day 0, duplicate direct-to-MGIT viability control tubes were set up by inoculating supplemented BACTEC MGIT tubes with the same volume of mycobacteria as the samples. The time to positivity (TTP) read-out was converted to log_10_ CFU using stock standard curves of TTP against inoculum volume and CFU. Results are presented as normalised BCG growth (log_10_ CFU of sample -log_10_ CFU of growth control).

### 2.5 Statistical analysis

Following normality testing (Shapiro-Wilk), longitudinal data was analysed using a one-way ANOVA with Dunnett’s correction for multiple comparisons (for normally-distributed data) or a Friedman test with Dunn’s correction for multiple comparisons (for non-normally distributed data). For studies with random missing values due to sample unavailability (Figure 1), data was logged and analysed using a mixed-effects model with Dunnett’s correction for multiple comparisons. When comparing two conditions (eg. serum vs. plasma), a paired t-test (for normally-distributed data) or Wilcoxon signed-rank test (for non-normally distributed data) was conducted. Associations were determined using a Spearman’s rank correlation.

## 3.0 Results

### 3.1 BCG vaccination in humans and macaques induces higher levels of PPD-specific antibodies compared with baseline

Using samples taken from UK adults enrolled into human Study 1, we observed a significant increase in PPD-specific IgG at 28 and 84 days post-BCG vaccination in serum (p=0.01 and p=0.0001 respectively, Figure 1A) and plasma (p=0.0004 and p=0.0002 respectively, Figure 1B) compared with baseline. PPD-specific IgA increased at 28 days post-BCG vaccination in serum (p=0.003, Figure 1D) and at days 28 and 84 in plasma (p=0.004 and p=0.0005 respectively, Figure 1E). As IgM is the first isotype to be produced following B cell stimulation, PPD-specific IgM was measured at earlier time-points (7, 14 and 21 days post-BCG vaccination), and we saw a significant increase at 14 and 21 days in plasma only (p=0.001 and p=0.0001 respectively, Figure 1G).

Similarly, in rhesus macaques enrolled into macaque Study 1, we observed a significant increase in PPD-specific IgG at 56 days post-BCG vaccination in serum and plasma (p=0.01 and p=0.004 respectively, Figure 2A-B) compared with baseline. PPD-specific IgA increased at 28 and 56 days post-BCG vaccination in plasma only (p=0.02 and p=0.02 respectively, Figure 2E), although there was a similar non-significant trend in serum (Figure 2D). There was a trend towards increased PPD-specific IgM taken at the same time-points but this was not significant; earlier time-points were not available for this study. No differences were observed over time in the control group for any of the isotypes measured for either species. There was no difference in fold change in PPD-specific IgG or IgA levels at 28 days following BCG vaccination (the only directly comparable time-point) in serum or plasma between humans and macaques (Figure S1A-B). We observed several associations between fold change following BCG vaccination in levels of different PPD-specific isotypes in both humans and macaques, as summarised in Tables S1–S4.

### 3.2 PPD-specific antibody levels, but not fold-change following vaccination, are higher in serum compared with plasma with a strong correlation between responses

In humans, we observed significantly higher levels of PPD-specific IgG in serum compared with plasma at all time-points (p=0.02, p=0.04 and p=0.007 at baseline, day 28 and day 84 respectively, Figure 1C). There were also higher levels of PPD-specific IgA in serum compared with plasma at all time-points measured (p=0.002, p=0.004 and p=0.01 at baseline, day 28 and day 84 respectively, Figure 1F), and higher levels of PPD-specific IgM in serum compared with plasma at all time-points measured (p=0.007, p=0.0001, p=0.005 and p=0.003 at baseline, day 7, day 14 and day 28 respectively, Figure 1I). Similarly, in macaques, there were significantly higher levels of PPD-specific IgG in serum compared with plasma at all time-points measured (p=0.0009, p=0.02 and p=0.04 at baseline, day 28 and day 56 respectively, Figure 2C), although only at 28 days for IgA (p=0.04), and there were no significant differences at any time-point for IgM (Figure 2I).

There was no difference in fold change following BCG vaccination between serum and plasma for any of the isotypes at any of the time-points measured in either humans (Figure S2A-C) or macaques (Figure S2D-F). The correlation between PPD-specific antibodies in the BCG vaccinated group in serum and plasma was strong at all time-points for IgG (p<0.0001 and r=0.97, 0.99 and 0.92 at baseline, day 28 and day 84; Figure 3A), IgA (p<0.0001 and r=0.88, 0.86 and 0.82 at baseline, day 28 and day 84; Figure 3B), and IgM (p<0.0001 and r=0.97, 0.97, 0.86 and 0.90 at baseline, day 7, day 14 and day 21; Figure 3C) in humans. Similarly, in macaques, the correlation between PPD-specific antibodies in serum and plasma was strong at all time-points for IgG (p=0.0001, 0.0005 and 0.002 and r=0.95, 0.92 and 0.88 at baseline, day 28 and day 56; Figure 3D), IgA (p=0.005, 0.002 and 0.0001 and r=0.83, 0.87 and 0.95 at baseline, day 28 and day 56; Figure 3E), and IgM (p=0.004, 0.0005 and <0.0001 and r=0.84, 0.92 and 0.99 at baseline, day 28 and day 56; Figure 3F).

### 3.3 Association between levels of serum antibodies specific to different mycobacterial fractions

Using serum collected from rhesus macaques enrolled into macaque Study 1, IgG, IgA and IgM responses to *M.tb* whole cell lysate (MTB WCL) were measured. There was a significant increase in MTB WCL-specific IgG at 28 and 56 days post-BCG vaccination (p=0.03 and p=0.03 respectively, Figure S3A) compared with baseline. MTB WCL-specific IgA increased at 56 days post-BCG vaccination in serum (p=0.01, Figure S3B). No differences were observed over time in MTB WCL-specific IgM (Figure S3C) or in the control group for any of the isotypes measured. There were significant associations between PPD-specific and MTB WCL-specific IgG at baseline and day 56 post-BCG vaccination (p=0.008 and p=0.03 respectively, Figure S3D), IgA at day 28 (p=0.001, Figure S3E) and IgM at day 28 (p=0.03, Figure S3F).

IgG levels specific to PPD and whole BCG were measured in serum taken at baseline and 28 days in human Study 1. Following BCG vaccination, there was a significant increase in IgG specific to whole BCG and PPD (p=0.006 and p=0.04 respectively, paired t-test, Figure S4A-B). There was no change in the control group over time. There was a significant correlation between BCG- and PPD-specific IgG levels at 28 days post-vaccination (p=0.005, Spearman’s correlation, Figure S4C), but not at baseline. Similarly, IgG levels specific to PPD and whole BCG were measured in serum taken at baseline and 56 days in macaque Study 2. There was a significant increase in both BCG-specific and PPD-specific IgG following BCG vaccination (p=0.04 and p=0.04 respectively, paired t-test, Figure S4D-E). There was a significant correlation between the two measures at baseline (p=0.04, Spearman’s correlation, Figure S4F). There was no change in PPD-specific or BCG-specific IgG levels in the control group over time in either study.

Using serum collected from rhesus macaques (macaque Studies 3 and 4) and cynomologus macaques (macaque Study 5), we compared responses between the two species at the same time-points following BCG vaccination with the standard dose. We observed similarly significant induction of PPD-specific IgG at 56 and 140 days post-BCG vaccination in the two species (p=0.0003 and p=0.001 respectively in rhesus macaques; p=0.001 and p=0.0007 respectively in Mauritian cynomolgus macaques; RM ANOVA with Dunnett’s post-test; Figure S5A). There was no difference in fold change in IgG response following BCG vaccination between the species at either time-point (Figure S5B).

In serum samples collected from Nepalese military recruits enrolled in human Study 2, IgG responses specific to multiple mycobacterial fractions were measured at baseline, 7 and 70 days later. At 70 days post-BCG vaccination, there was a significant increase in IgG specific for whole BCG, whole-cell lysate, cell membrane fraction, culture filtrate and lipoarabinomannan (LAM) compared with baseline (p=0.005, p=0.01, p=0.002, p=0.01 and p=0.02 respectively, RM ANOVA with Dunnett’s post-test, Figure 4A-E). There was no change over time in the level of IgG specific to any of the antigens in the historically BCG-vaccinated group who received no intervention. The greatest BCG-induced fold change was in IgG specific for *M.tb* whole cell lysate and *M.tb* culture filtrate (median fold change = 1.7 and 1.6 respectively), while the smallest was against *M.tb* cell membrane fraction and LAM (median fold change = 1.2 and 1.2). However, there was a significant association in BCG-induced fold change in IgG between all mycobacterial fractions (Table 2). There was no difference in fold change in the BCG-specific IgG response following BCG vaccination between individuals in Study 1 (from a non-TB endemic country) at 84 days and individuals from Study 2 (from a TB-endemic country) at 70 days (Figure S1C).

**Table 2.** Spearman’s correlations between fold change (post-vaccination/baseline) in IgG specific to different mycobacterial fractions following BCG vaccination in serum collected from Nepalese military recruits.

|  | Whole BCG | M.tb Whole cell lysate | M.tb cell membrane fraction | M.tb culture filtrate | LAM |
| --- | --- | --- | --- | --- | --- |
| Whole BCG |  | .955<br>.000<br>*** | .791<br>.004<br>** | .909<br>.000<br>*** | .809<br>.003<br>** |
| M.tb whole cell lysate | .955<br>.000<br>*** |  | .745<br>.008<br>** | .964<br>.000<br>*** | .718<br>.013<br>* |
| M.tb cell membrane fraction | .791<br>.004<br>** | .745<br>.008<br>** |  | .782<br>.004<br>** | .927<br>.000<br>*** |
| M.tb culture filtrate | .909<br>.000<br>*** | .964<br>.000<br>*** | .782<br>.004<br>** |  | .736<br>.010<br>** |
| LAM | .809<br>.003<br>** | .718<br>.013<br>* | .927<br>.000<br>*** | .736<br>.010<br>** |  |

### 3.4 BCG-specific IgG-secreting plasmablasts and memory B cells are induced following BCG vaccination

To investigate the origin of the antibodies observed, B cell ELISpot assays were developed to detect BCG-specific IgG-secreting cells indicative of either plasmablasts measured following an overnight assay, or memory B cells (mBCs) enumerated following several days of culture and stimulation with polyclonal mitogens. The antigen used was the same BCG strain given during vaccination (BCG-SSI). The transient induction of plasmablasts was noted at 7 days following BCG vaccination in Group A (mean of 20 ASC per million PBMC or 1.7% of total IgG+ ASC; p=0.0009), which was undetectable by day 70 (Figure 5A-B). No plasmablast responses were seen in Group B (historically BCG vaccinated but receiving no intervention). There was an increase in the memory B cell response at 7 and 70 days post-BCG vaccination in Group A (p=0.03 and p<0.0001 respectively), and Group B showed persisting stable BCG-specific mBC responses (mean of 16 mBC-derived ASC per million cultured PBMC or 0.06% of IgG+ mBC-derived ASC) (Figure 5C-D), reflecting their historical BCG vaccination status. There was a trend towards a correlation between ASC responses at 7 days and BCG-specific IgG levels at 7 days (Figure 5E), which was significant at 70 days (p=0.003, r=0.83, Spearman’s rank correlation, Figure 5F). There were no associations between mBC-derived ASC responses and BCG-specific IgG levels at either time-point studied.

**Figure 5.**
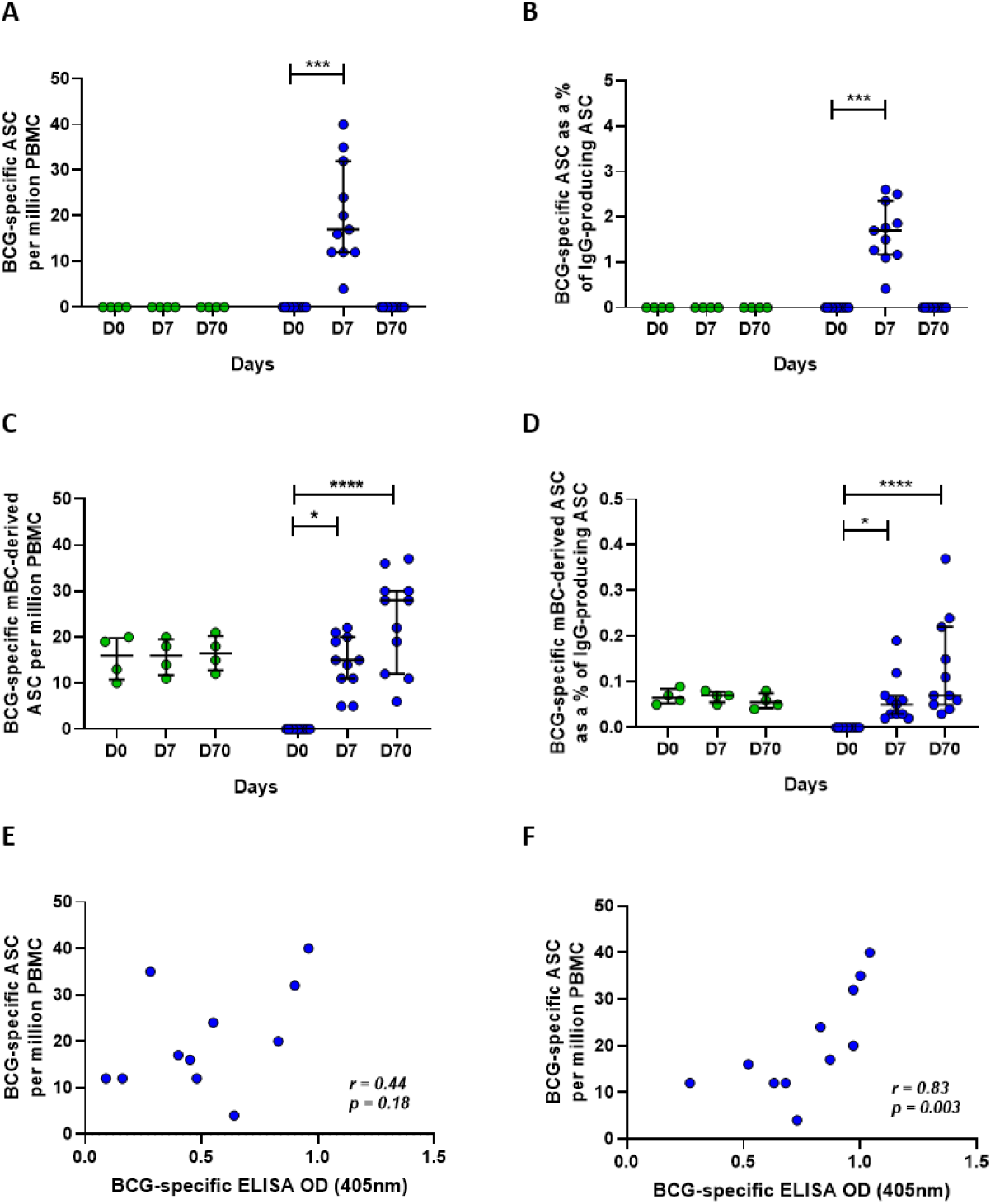
BCG-specific IgG-secreting plasmablast (ASC) and memory B cell-derived ASC responses following BCG vaccination. ASC and mBC-derived ASC responses were determined at baseline, 7 days and 70 days using cells collected from Nepalese military recruits enrolled into human Study 2 (green = historically BCG-vaccinated individuals who received no intervention, blue = BCG-naïve individuals who received BCG vaccination). ASC responses are presented as the number of BCG-specific ASCs per million PBMC (A) and as the proportion (%) of total IgG-secreting ASC (B). mBC responses are shown as the number of BCG-specific mBC-derived ASC per million cultured PBMC (C) and as the proportion (%) of mBC-derived ASC of total IgG-secreting ASC (D). Points represent the mean of triplicates at two dilutions and bars represent the median with IQR. A Friedman test with Dunn’s multiple comparisons test, where * indicates a p-value of <0.05, ** indicates a p-value of <0.01, *** indicates a p-value of <0.001 and **** indicates a p-value of <0.0001. A Spearman rank correlation was performed between ASC responses and BCG-specific IgG responses measured at 7 days (E) and 70 days (F) after BCG vaccination.

### 3.5 Antibodies may play a functional role in the control of mycobacterial growth *in vitro*

The direct PBMC MGIA was conducted using cryopreserved cells and autologous serum collected at baseline and 84 days post-BCG vaccination from a subset of volunteers, n=9, enrolled into human Study 1. When serum was matched to time-point, there was a significant improvement in control of mycobacterial growth at 84 days post-BCG vaccination compared with baseline, as previously reported [45] (p=0.02, Figure 6A). When serum was swapped between pre- and post-vaccination time-points (baseline PBMC cultured with day 84 serum, and day 84 PBMC cultured with baseline serum), control of mycobacterial growth was no longer different between time-points. This finding was validated using samples (n=9) collected at baseline and 70 days taken from Study 2. When serum was matched to time-point, there was a significant improvement in control of mycobacterial growth at 70 days post-BCG vaccination compared with baseline (p=0.0007, Figure 6B). When serum was exchanged between pre- and post-vaccination time-points (baseline PBMC cultured with autologous day 70 serum, and day 70 PBMC cultured with autologous baseline serum), the difference in control of mycobacterial growth was lost. Finally, control of mycobacterial growth was modestly improved using cells taken at baseline cultured with autologous serum taken at 70 days post-vaccination compared with autologous serum taken at baseline (p=0.03, Figure 6B).

**Figure 6.**
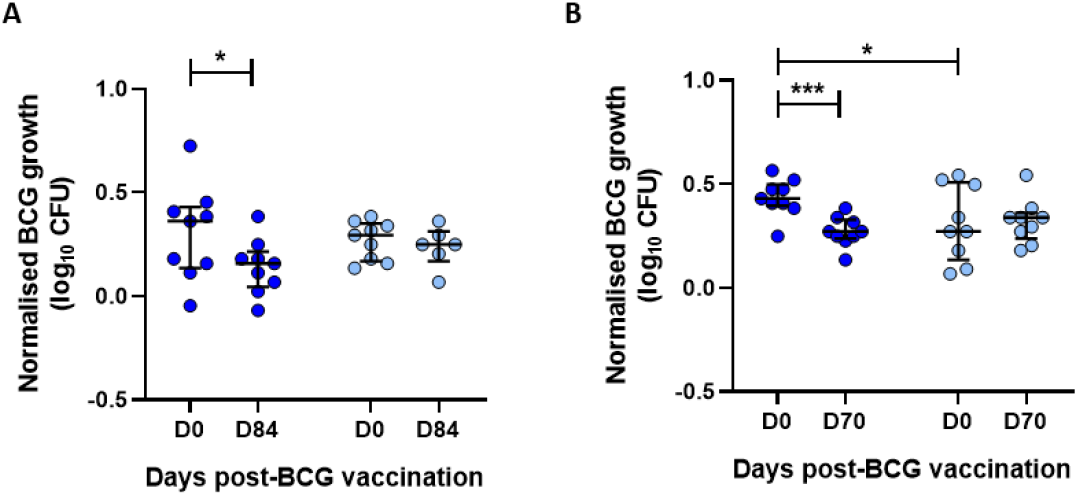
Contribution of autologous time-point matched serum to BCG vaccine-induced control of mycobacterial growth. Samples were used from n=9 healthy UK adults enrolled into human Study 1 (A) and n=9 healthy Nepalese military recruits enrolled into human Study 2 (B), all of whom received ID BCG vaccination. The direct MGIA was performed using PBMC collected at baseline and 84 days (A) or 70 days (B) post-BCG vaccination, using either autologous serum matched to time-point (dark blue), or autologous serum switched by time-point ie. post-vaccination serum cultured with baseline PBMC and baseline serum cultured with post-vaccination PBMC (light blue). Points represent the mean of single or duplicate cultures and bars show the median with the interquartile range (IQR). A paired t-test was used to compare between time-points, where * indicates a p-value of <0.05, ** indicates a p-value of <0.01.

## 4.0 Discussion

We have observed significant induction of specific IgG following BCG vaccination in two independent human cohorts (one from a TB-endemic region and one from a non-endemic region). Our findings are consistent with several previous reports [46–49]. While others observed no change in IgG titre following BCG vaccination, limitations of such studies have included antibody quantification methods, small sample sizes, variable/unspecified time-points or duration of follow-up, differences in dose and strain of BCG, and variable baseline characteristics of the study populations [26, 50–52]. Both of our cohorts comprised healthy adults who received vaccination with a standard dose of BCG SSI and antibody levels were measured up to ~3 months post-vaccination. Interestingly, in a comparison of different antigen fractions, the greatest fold change was in IgG specific for *M.tb* whole cell lysate and culture filtrate, suggesting recognition of secreted as well as surface proteins. The strong associations between responses to whole BCG and different *M.tb* fractions is reflective of the high degree of homology (>95%) between the species, and supports the induction of responses relevant to the target pathogen [53].

While macaques are partially protected by BCG vaccination and considered the most relevant preclinical model for human TB vaccine development, there is a paucity of literature on the antibody response to BCG in this model [54–56]. We show significant BCG vaccine-mediated induction of specific IgG consistently across five different macaque studies, which included two species and animals of different ages (young adults aged 2.4-4.2 years and older adults aged 14-15 years). Interestingly, IgG and IgA levels were increased at 56 days; contrary to a recent report in the same species that suggested a peak in both isotypes at 28 days post-vaccination with BCG administered by different routes (including ID) [16]. The lesser IgM response could indicate that the IgG and IgA responses represent secondary memory responses, although at the relatively late time-points available for NHP, B cells may have already undergone class-switching.

We saw similar levels of IgG in rhesus and cynomolgus macaques with a similar fold change in titre relative to baseline in both species; whereas a previous study reported a weak *M.tb*-specific IgG response in rhesus but not cynomolgus macaques at 14 weeks post-vaccination, it should be noted that animals received 3 weeks of daily oral isoniazid/rifadin at 8 weeks [33]. Rhesus macaques are more susceptible to acute progressive TB disease and BCG confers lower efficacy compared with cynomolgus macaques [33, 57]; our finding of similar total specific IgG responses between the species may therefore suggest that these antibodies are not protective, or that different attributes (such as antigen-specificity, subtype, post-translational modifications, affinity or avidity) are more relevant. Importantly, we observed similar fold change in IgG and IgA levels relative to baseline in humans and macaques at the comparable time-point (day 28), offering some confidence in the translatability of this model in this context.

Serum is often a limited resource due to restricted blood collection volumes. We therefore assessed whether plasma, a by-product of routine cell separation, represents an equivalent alternative in the context of measuring BCG vaccine-induced specific antibody responses. Our finding that PPD-specific antibody levels in serum and plasma correlate strongly is consistent with the observations of Siev et al., who concluded that serum and plasma can be used interchangeably to test for antibody responses to mycobacterial antigens even in the same assay [58]. However, we observed PPD-specific antibody responses that were modestly but significantly higher in serum than plasma across isotypes and species. Other studies have similarly noted lower levels of proteins in plasma than serum [59, 60], potentially associated with the addition of anticoagulant (such as heparin) to blood prior to obtaining plasma, or the presence of fibrinogen which may influence antibody-antigen binding [61]. However, fold change in BCG vaccine-induced antibody responses was comparable between serum and plasma indicating that sensitivity to detect a vaccine-induced response does not differ.

To our knowledge, the specific antibody-secreting B cell response to BCG vaccination has not been previously described. Using an overnight ELISpot assay, we observed induction of BCG-specific IgG-secreting plasmablasts at 7 days post-BCG vaccination, returning to baseline by day 70. This is consistent with the kinetic of response to other primary vaccinations including those against rabies and influenza [38, 39]. The poor correlation between ASC responses and early IgG antibody induction is perhaps to be expected; at 7 days following vaccination, BCG-specific IgG would still be in the very early stages of production. While a significant association was noted between ASC responses and IgG levels at 70 days post-vaccination, it may be that levels have started to wane by this time and that determining responses at an earlier time-point would provide an even stronger association. Strong correlations of IgG plasma cell responses with IgG antibody responses after primary rabies immunisation by area under the curve analysis has been reported elsewhere, but our limited time-points did not permit such analysis [38].

BCG-specific mBC responses were detectable in the historically BCG-vaccinated group, who had received BCG up to 20 years previously, at all time-points. This is in concordance with a previous study of historic BCG vaccinees assessed 13-45 years post-vaccination [41]. The responses noted here are higher than previously seen, which may be due to the use of different antigens (whole BCG vs. PPD) and different study populations. In the previously BCG-naïve group, mBCs induced by BCG vaccination were apparent at 7 days and had further increased by 70 days, consistent with the detection of mBCs in peripheral blood by 7 days post-immunisation in a murine model [62]. A study assessing longitudinal mBC responses following exposure to candidate malaria vaccines showed a gradual increase in specific mBC after immunisation which peaked at 84 days before declining by 140 days [40]. One limitation of our study is the inability to define the peak or longevity of mBC responses due to logistical restrictions on collecting later follow-up samples, but one would similarly expect contraction to a persisting residual level over time [63].

Interestingly, we saw relatively high antibody responses at baseline and in the naïve unvaccinated controls across isotypes and studies, which may indicate cross-reactive antibodies induced by environmental exposure to non-tuberculous mycobacteria (NTM) [64, 65]. Others have also noted pre-existing specific antibodies in serum from PPD-negative individuals with no known exposure to *M.tb* [27, 66]. While NTM are prevalent in Nepal, Western countries are generally considered to have low circulating levels, but there are indications of a steep rise in incidence in the UK in recent decades [67, 68]. The macaques included here were housed partially outdoors, and as such may also have had environmental exposure. It is possible that an element of the ASC response observed in the Nepalese cohort was due to the generation of plasmablasts from pre-existing mycobacteria-specific mBCs. Indeed, the rapidity and magnitude of the ASC response was comparable to recall responses observed in secondary vaccination studies [38, 40]; although this hypothesis is not supported by the lack of mBC responses at baseline in unvaccinated volunteers, they may have been present at frequencies below the limit of detection of the ELISpot assay. It remains to be determined whether baseline reactivity interferes with induction of a humoral response to BCG vaccination in the way it appears to affect IFN-γ responses [69], particularly given the widely-accepted masking/blocking hypotheses for the geographical variation in BCG efficacy [12].

In order to explore a potential functional role for the antibodies detected, we applied an unbiased sum-of-the-parts mycobacterial growth inhibition assay (MGIA) that we have previously optimised, standardised and harmonised as a potential surrogate of TB vaccine-induced protection [45, 70]. Using this assay, we (and others) have demonstrated ability to detect a BCG vaccine-induced response and an association between the outcome of the direct MGIA and protection from *in vivo* mycobacterial infection [44, 71–74]. Various immune mechanisms have been proposed to contribute to mycobacterial control in the direct MGIA including polyfunctional CD4+ T cells and trained innate immunity [44, 72, 73]. In two independent cohorts, we confirmed previous findings of enhanced control of mycobacterial growth in the direct MGIA following BCG vaccination, and importantly showed that this effect was cancelled out by exchanging pre-for post-vaccination serum and vice versa.

Further work is required to determine the serum component/s responsible, but other studies have reported that post-BCG vaccination serum can enhance the control of mycobacterial growth *in vitro* via antibody-mediated mechanisms [27, 28], and we hypothesise that antibodies are likewise playing a role in our assay. Indeed, we have previously shown that IgG1 responses to *M.tb-specific* antigens correlated with improved mycobacterial control in a study of individuals with ATB or LTBI [43], and that specific IgG responses following *in vivo* mycobacterial infection in historically BCG vaccinated volunteers were associated with control of mycobacterial growth following *in vitro* infection in the direct MGIA [44]. A role for antibodies may explain in part why we saw no association between outcome of the direct whole blood and PBMC MGIAs when pooled human serum rather than autologous serum was used in the latter [75]. In our second human cohort, we observed enhanced mycobacterial control when baseline cells were cultured with post-vaccination serum compared with baseline cells cultured with baseline serum, although the effect was less pronounced than between baseline cells cultured with baseline serum and post-vaccination cells cultured with post-vaccination serum. This suggests that control of mycobacterial growth in the direct MGIA is mediated by a combination of cellular and humoral immunity, and supports the recommendation to use autologous serum in the assay as a more representative *ex vivo* approach.

In conclusion, we have demonstrated BCG vaccine-mediated induction of antibodies specific to PPD and whole BCG that are modestly higher in serum than plasma and comparable across humans and macaques, as well as serum responses to a range of mycobacterial fractions in humans. We have also shown for the first time the rapid and transient induction of antibody-secreting plasmablasts following BCG vaccination, together with a robust memory B cell response at 70 days post-vaccination. While contracted, a measurable mBC response was still present in volunteers that received BCG vaccination up to 20 years previously, suggesting that this pool is maintained and may contribute to a long-lived humoral response. Finally, we used a functional *in vitro* MGIA to demonstrate a potential contribution of the antibody response to BCG vaccine-mediated control of mycobacterial growth. Our findings indicate that the humoral immune response in the context of BCG vaccination merits further attention, particularly as the majority of TB vaccine candidates are intended as a boost to BCG. Further characterisation of the BCG-induced response may aid in the design of more efficacious TB vaccine candidates that could benefit from the induction of humoral as well as cellular immunity.

## 5.0 Tables

**Table S1.** Spearman’s correlations between fold change (post-vaccination/baseline) in different isotypes following BCG vaccination in serum collected from healthy UK adults.

|  |  | IgG |  | IgA |  | IgM |  |  |
| --- | --- | --- | --- | --- | --- | --- | --- | --- |
|  |  | D28 FC | D84 FC | D28 FC | D84 FC | D7 FC | D14 FC | D21 FC |
| IgG | D28 FC |  | .846<br>.000<br>*** | .667<br>.003<br>** | .824<br>.001<br>** | .279<br>.315 | .527<br>.096 | .358<br>.310 |
|  | D84 FC | .846<br>.000<br>*** |  | .495<br>.086 | .697<br>.006<br>** | -.005<br>.986 | .286<br>.535 | .750<br>.052 |
| IgA | D28 FC | .667<br>.003<br>** | .495<br>.086 |  | .609<br>.021<br>* | .015<br>.958 | .373<br>.259 | .248<br>.489 |
|  | D84 FC | .824<br>.001<br>** | .697<br>.006<br>** | .609<br>.021<br>* |  | .308<br>.331 | .679<br>.094 | .383<br>.383 |
| IgM | D7 FC | .279<br>.315 | -.005<br>.986 | .015<br>.958 | .308<br>.331 |  | .588<br>.074 | -.450<br>.224 |
|  | D14 FC | .527<br>.096 | .286<br>.535 | .373<br>.259 | .679<br>.094 | .588<br>.074 |  | .383<br>.308 |
|  | D21 FC | .358<br>.310 | .750<br>.052 | .248<br>.489 | .383<br>.383 | -.450<br>.224 | .383<br>.308 |  |

**Table S2.** Spearman’s correlations between fold change (post-vaccination/baseline) in different isotypes following BCG vaccination in serum collected from rhesus macaques.

|  |  | IgG |  | IgA |  | IgM |  |
| --- | --- | --- | --- | --- | --- | --- | --- |
|  |  | D28 FC | D56 FC | D28 FC | D56 FC | D28 FC | D56 FC |
| IgG | D28 FC |  | .648<br>.043<br>* | .903<br>.000<br>*** | .661<br>.038<br>* | .697<br>.025<br>* | .709<br>.022<br>* |
|  | D56 FC | .648<br>.043<br>* |  | .552<br>.098 | .988<br>.000<br>** | .600<br>.067 | .758<br>.011<br>* |
| IgA | D28 FC | .903<br>.000<br>*** | .552<br>.098 |  | .564<br>.090 | .648<br>.043<br>* | .503<br>.138 |
|  | D56 FC | .661<br>.038<br>* | .988<br>.000<br>** | .564<br>.090 |  | .539<br>.108 | .782<br>.008<br>** |
| IgM | D28 FC | .697<br>.025<br>* | .600<br>.067 | .648<br>.043<br>* | .539<br>.108 |  | .600<br>.067 |
|  | D56 FC | .709<br>.022<br>* | .758<br>.011<br>* | .503<br>.138 | .782<br>.008<br>** | .600<br>.067 |  |

**Table S3.**
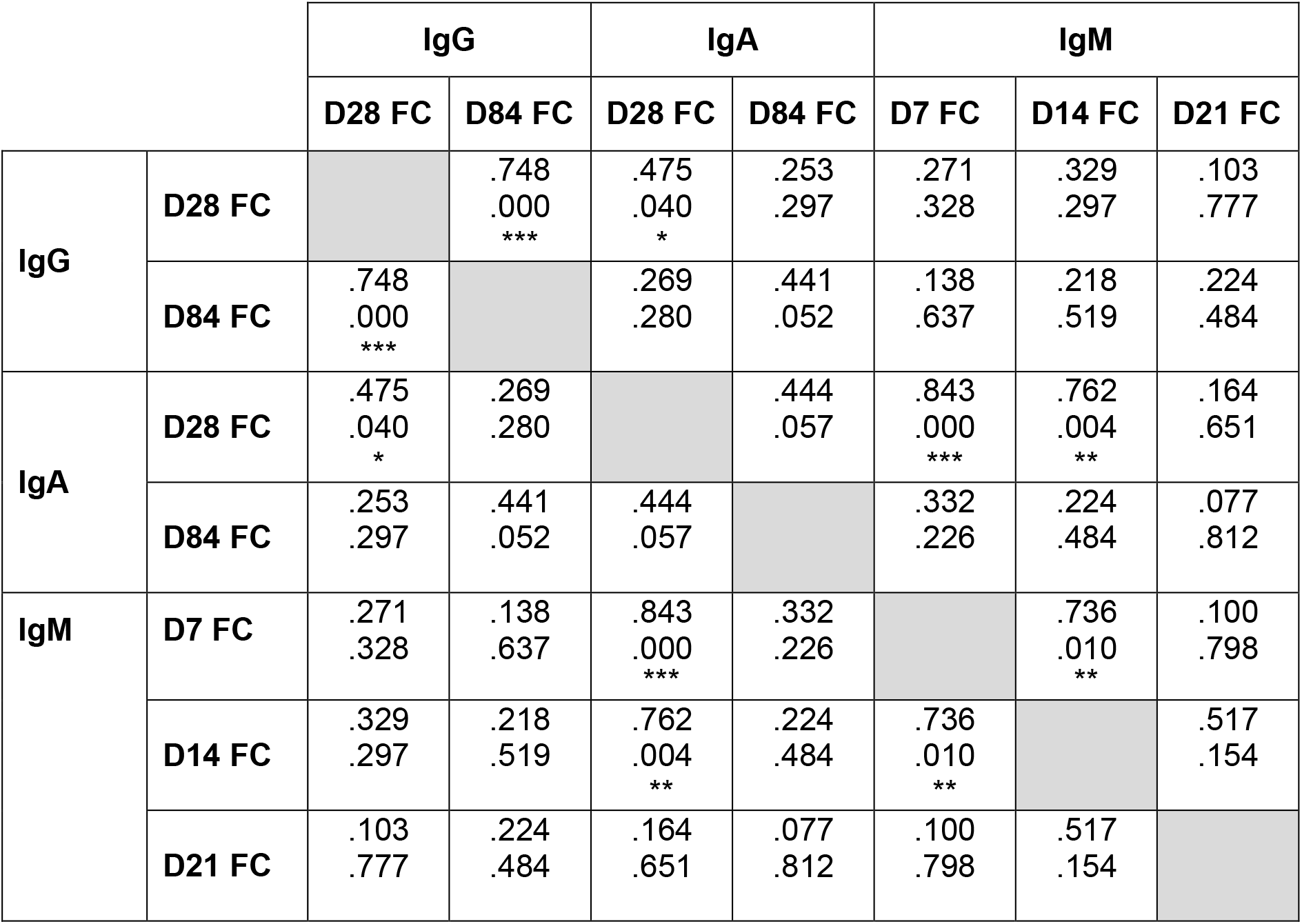
Spearman’s correlations between fold change (post-vaccination/baseline) in different isotypes following BCG vaccination in plasma collected from healthy UK adults.

**Table S4.** Spearman’s correlations between fold change (post-vaccination/baseline) in different isotypes following BCG vaccination in plasma collected from rhesus macaques.

|  |  | **IgG** |  | **IgA** |  | **IgM** |  |
| --- | --- | --- | --- | --- | --- | --- | --- |
|  |  | **D28 FC** | **D56 FC** | **D28 FC** | **D56 FC** | **D28 FC** | **D56 FC** |
| **IgG** | **D28 FC** |  | .709<br>.022<br>* | .733<br>.016<br>* | .564<br>.090 | .842<br>.002<br>** | .382<br>.276 |
|  | **D56 FC** | .709<br>.022<br>* |  | .394<br>.260 | .903<br>.000<br>*** | .479<br>.162 | .418<br>.229 |
| **IgA** | **D28 FC** | .733<br>.016<br>* | .394<br>.260 |  | .394<br>.260 | .648<br>.043<br>* | .576<br>.082 |
|  | **D56 FC** | .564<br>.090 | .903<br>.000<br>*** | .394<br>.260 |  | .261<br>.467 | .539<br>.108 |
| **IgM** | **D28 FC** | .842<br>.002<br>** | .479<br>.162 | .648<br>.043<br>* | .261<br>.467 |  | .467<br>.174 |
|  | **D56 FC** | .382<br>.276 | .418<br>.229 | .576<br>.082 | .539<br>.108 | .467<br>.174 |  |

## 7.0 Acknowledgements

We would like to thank all of the study volunteers, the clinical teams in Oxford and Birmingham for recruitment and assistance with Human Study 1, and Gurkha Brigade Headquarters and the clinical staff at Infantry Training Centre (ITC) Catterick, particularly Donna Tupper, for their support and assistance with Human Study 2. We thank the staff of the Biological Investigations Group at PHE Porton Down and the PHE macaque colonies who assisted in conducting the macaque studies. We also thank Victoria Harris (Primary Care Health Sciences, University of Oxford) for advice on some aspects of the statistical analysis.

## 8.0 Author contributions

RT, MO and HM conceived and designed the work; JB, MO, RT, MPPA, MW, AJ, SH, DW, SS, SE, AW and SGS conducted the studies and/or contributed to the acquisition of data; RT, JB, MO and HM contributed to the interpretation of the data. RT, JB and MO wrote the paper and all authors provided critical analysis and approved the submitted version.

---

**Figure S1.**
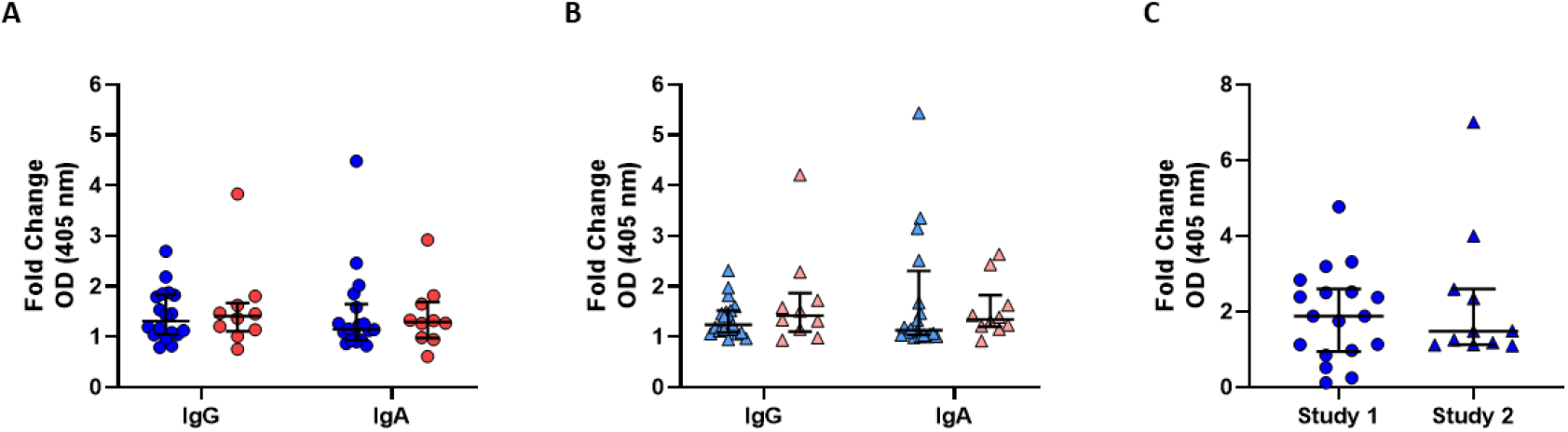
Comparison of PPD-specific IgG responses to BCG vaccination between humans and macaques and between different human populations. Fold change (28 days post-BCG vaccination/baseline) in PPD-specific IgG in serum (circles, A) and plasma (triangles, B) was compared between healthy UK adults enrolled into human Study 1 (blue) and rhesus macaques enrolled into macaque Study 1 (red). Fold change in PPD-specific IgG following BCG vaccination was compared between adults from the UK (non-TB endemic) enrolled in human Study 1 at day 84 (circles) and adults from Nepal (TB-endemic) human Study 2 at day 70 (triangles) (C). Points represent the mean of triplicate values and bars show the median with the interquartile range (IQR).

**Figure S2.**
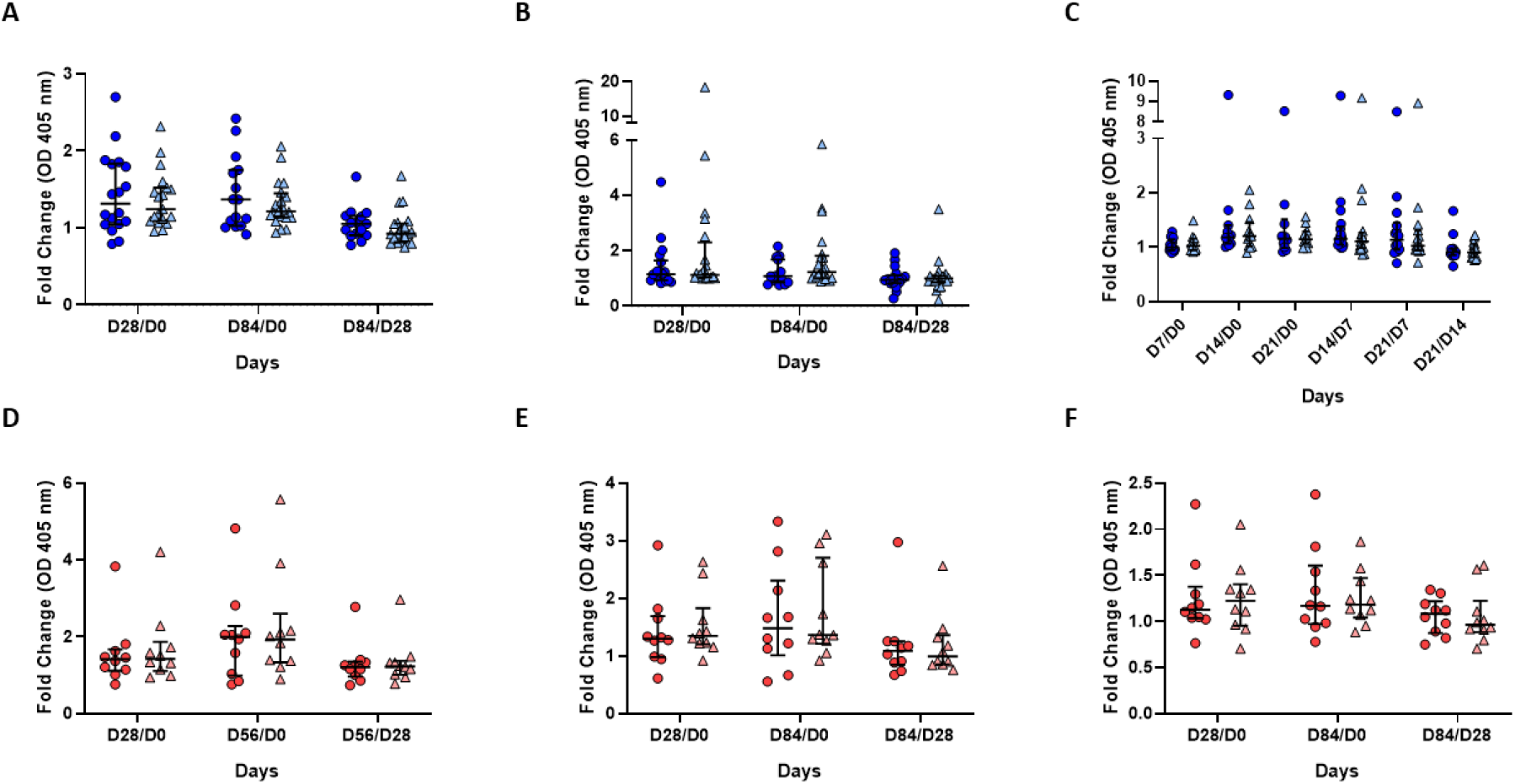
Comparison of fold change in PPD-specific antibody levels following BCG vaccination between serum and plasma. Fold change in PPD-specific antibody responses following BCG vaccination was compared between serum (circles) and plasma (triangles) collected from healthy UK adults enrolled into human Study 1 (blue, A-C) and rhesus macaques enrolled into macaque Study 1 (red, D-F). Fold change in IgG (A), IgA (B) and IgM (C) responses were compared between baseline and 28 days post-vaccination, baseline and 84 days post-vaccination, and 84 and 28 days post-vaccination in humans. Fold change in IgG (D), IgA (E) and IgM (F) responses were compared between baseline and 28 days post-vaccination, baseline and 56 days post-vaccination, and 56 and 28 days post-vaccination in macaques. Points represent the mean of triplicate values and bars show the median with the interquartile range (IQR).

**Figure S3.**
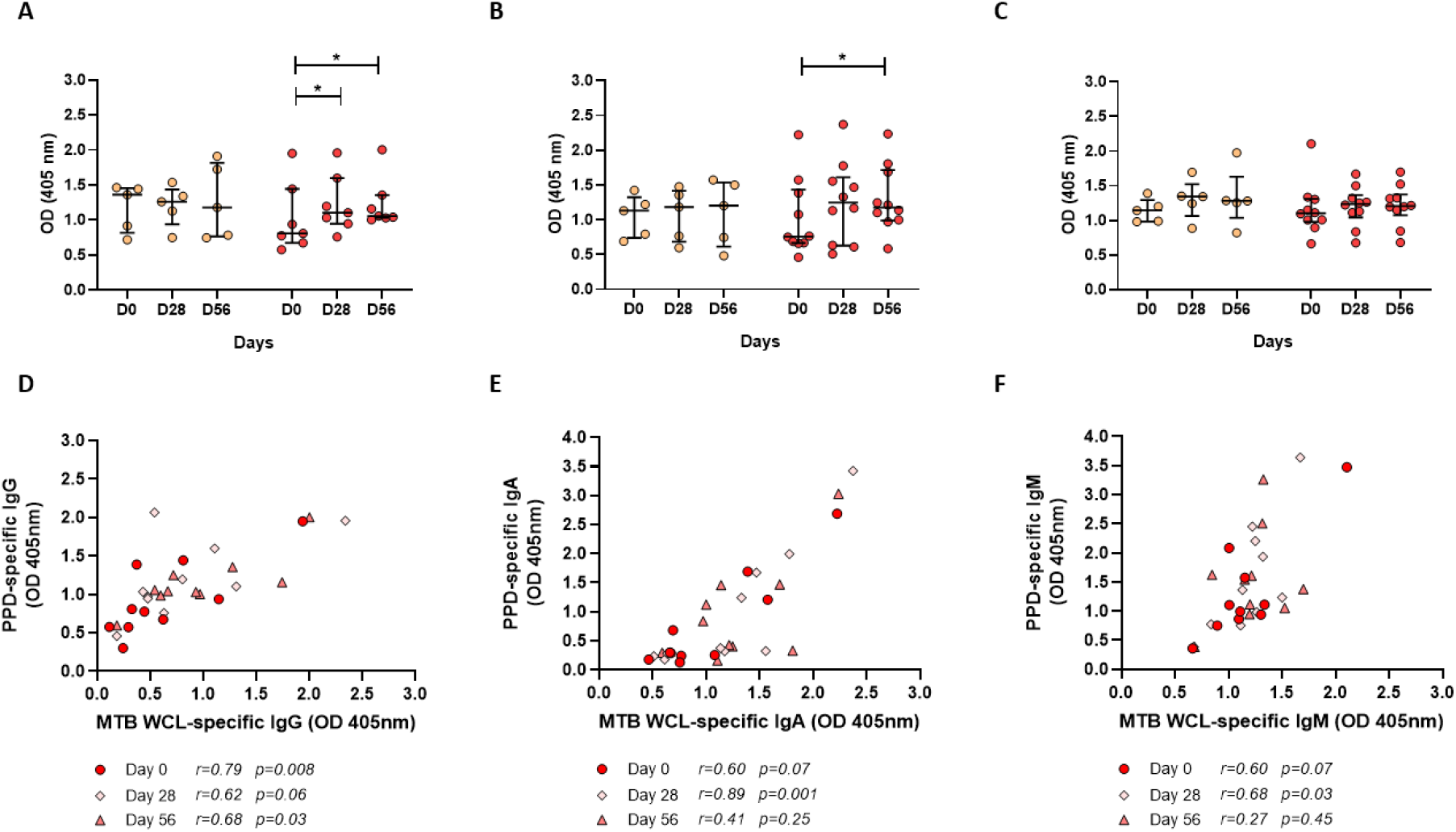
MTB WCL-specific antibody responses to BCG vaccination in rhesus macaques. Serum was collected from animals enrolled into macaque Study 1 which were either unvaccinated controls (orange) or received BCG vaccination (red). MTB WCL-specific IgG (A), IgA (B) and IgM (C) responses were measured in serum over time, and associations determined in responses following BCG vaccination to MTB WCL and PPD for IgG (D), IgA (E) and IgM (F). Points represent the mean of triplicate values and bars show the median with the interquartile range (IQR). A Friedman test with Dunn’s correction for multiple comparisons was used to compare the BCG vaccine-induced response between post-vaccination and baseline time-points (A-C), and a Spearman’s rank correlation was used to determine associations (D-F). * indicates a p-value of <0.05, ** indicates a p-value of <0.01, and *** indicates a p-value of <0.001.

**Figure S4.**
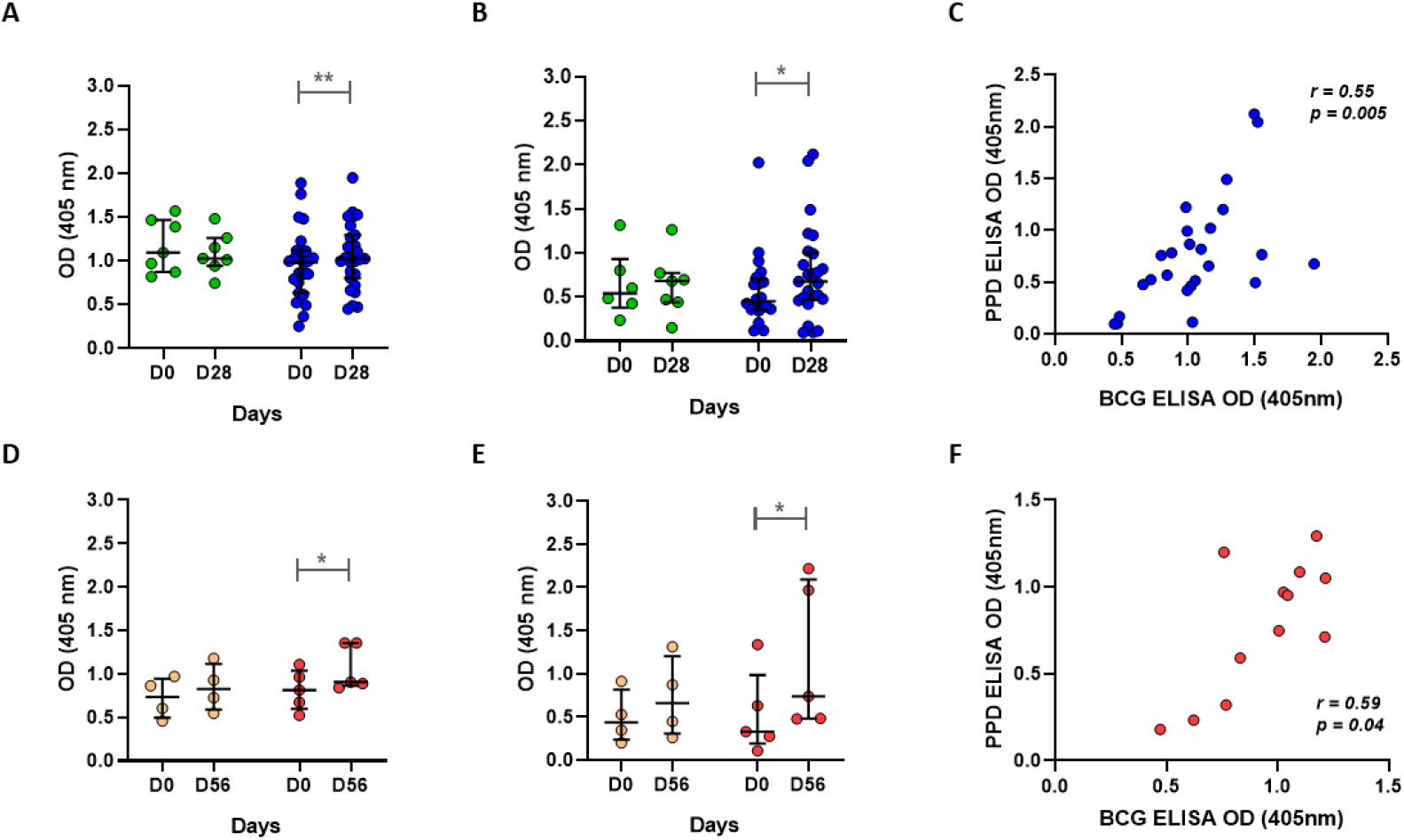
BCG-specific and PPD-specific IgG responses to BCG vaccination in serum. IgG responses in serum collected from healthy UK adults enrolled into human Study 1 (green = unvaccinated controls, blue = BCG vaccinees, A-C), rhesus macaques enrolled into macaque Study 3 (orange = unvaccinated controls, red = BCG vaccinees, D-F). IgG responses were compared to either whole BCG (A, D) or to PPD (B, E), and the association between the two was determined (C, F). Points represent the mean of triplicate values and bars show the median with the interquartile range (IQR). A Wilcoxon test was used to compare between time-points in the BCG vaccinated groups (A, B, D, E), and a Spearman’s rank correlation was used to determine associations. * indicates a p-value of <0.05, ** indicates a p-value of <0.01 and *** indicates a p-value of <0.001.

**Figure S5.**
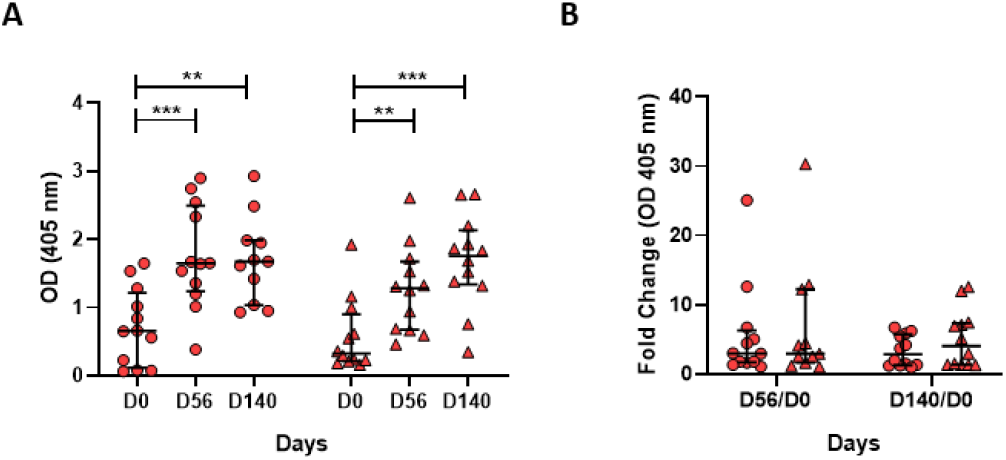
Comparison of specific IgG responses to BCG vaccination between rhesus and cynomolgus macaques. PPD-specific IgG levels (A) and fold change in IgG (B) following BCG vaccination were determined in rhesus macaques (circles) and Mauritian cynomolgus macaques (triangles) enrolled in macaque Study 4. Points represent the mean of triplicate values and bars show the median with the interquartile range (IQR). For A), an RM ANOVA with Dunnett’s multiple comparisons test was performed where * indicates a p-value of <0.05, ** indicates a p-value of <0.01 and *** indicates a p-value of <0.001.

